# Parents’ neural dynamics in response to their children’s speech

**DOI:** 10.64898/2026.09.25.754219

**Authors:** Steven L. Elmlinger, Ella Rosenberg, Sagi Jaffe-Dax, Mira L. Nencheva, Crystal Lee, Jessica E. Kosie, Casey Lew-Williams

## Abstract

The development of speech in human children is a protracted process that depends on social experience. From early life, young children produce vocalizations in the presence of others, which affords unique opportunities for feedback about how to communicate successfully. However, for feedback to happen, young children’s speech needs to effect change in the brains of their caregivers. Using dual-brain functional near-infrared spectroscopy (fNIRS) to measure real-time communication between caregivers and their 2-to 3-year-old children (*n* = 56), we found that children’s vocalizations selectively modulated caregivers’ brain activity. Parents’ brains showed a rapid response to children’s production of speech that was communicatively advanced (multiword utterances) vs. less advanced (babble). The findings imply that if part of learning to talk is figuring out how to get your communication partner to listen, then early conversations likely present continuous opportunities for feedback that guide children toward advances in spoken language.

## INTRODUCTION

What drives children to continue learning spoken language across time? Children could learn a small number of useful words and then stop. But that is not what children do—they work at learning to speak, incorporating speech sounds, words, and constructions into their repertoire for many years on end (McLeod and Crowe, 2018). Most research on this process focuses on caregiver speech directed to children (i.e., child-directed speech; (Weisleder and Fernald, 2013). However, when *children* produce speech and its precursors, they create moments rich with opportunity for social and language learning (Kuhl, 2007). Their voices change others’ behaviors (Elmlinger et al., 2025) and neural activity (Bornstein et al., 2017), the heart of communicative success (Nikolaus and Fourtassi, 2023). Thus, an important context of learning to speak is vocal exchange during social interactions (Romeo et al., 2021, 2018). However, little is known about how young children’s speech use shapes the real-time neural dynamics of interactions with caregivers.

Infants’ ability to shape ongoing social interaction with their voice facilitates communicative development. Prelinguistic infants use vocal behaviors to elicit feedback from their caregiver, guiding them to more advanced vocal patterns (Goldstein et al., 2009, 2003), and this social feedback system drives communicative advances across many species (Carouso-Peck and Goldstein, 2019; Takahashi et al., 2017; West and King, 1988). Once children incorporate words into their production, feedback from caregivers is sometimes thought of by researchers as negative in form—giving children a sense of when their communicative attempts go awry (Chouinard and Clark, 2003). Recently, however, a new form of positive feedback has been proposed—adults’ child-directed *listening*. The idea is that simply getting one’s communication partner to listen may constitute a feedback signal to the learner that guides them toward mature language use (Meylan et al., 2023; Newport et al., 1977; Nikolaus and Fourtassi, 2023). Adults’ listening to children talk seems to recruit an active prediction process to try to understand the unfolding utterances of their child (Meylan et al., 2023). When children’s early speech influences listeners, this communicative success itself may serve as a positive guiding cue to children about what to learn next. The only way for communicative success to occur is by influencing others’ brains.

What are the neural dynamics of adults’ child-directed listening? Adults’ and children’s brain activity tracks the ongoing behaviors of one another during natural interaction (Nguyen et al., 2020; Piazza et al., 2020). In order for child-directed listening to serve as feedback, the extent to which children’s vocalizations influence parents’ brain should, in aggregate, be sensitive to the communicative maturity of children’s utterances. At 2-3 years of age, children intermix reflexive sounds, babbling, and speech vocalizations (Vihman et al., 2009). Is there structure to how parents’ brains’ respond to different communicative behaviors from children? In order to examine variability in children’s learning trajectories, an important precondition is to be able to detect a neural signal of parents’ listening during real interactions. Here, we provide the first investigation of parents’ brain dynamics during real-time natural interaction as they listen to their own child’s vocalizations which vary in communicative complexity during natural interaction.

We examined a wide range of young children’s early vocalizations – spanning reflexive sounds, babbling, single words, and multiword utterances – and their influence on parents’ brain activity. This is a foundational step for understanding the puzzle of how children’s speech and parents’ responses become a feedback loop that furthers children’s learning over time. Listening to a child is a nonobvious, yet fundamental act in family life that may play an outsized role in children’s development. Listening to children may be a quite effortful process according to previous computational work (Meylan et al., 2023). If parents’ brains react similarly to any vocalization, then there would be little structure guiding children’s continued learning. However, if parents’ brains respond differently across the gradient of children’s communicative maturity, then that may signal to children what is of greater value in their communicative environment.

## RESULTS

### Child-directed listening in parents’ brains

To understand the temporal dynamics between children’s utterances and parents’ brain responses, we ran brain-behavior regressions by shifting the two signals relative to one another in time, taking regressions per dyad from –15 second lags (brain leading) to +15 s lags (child utterance leading) in 1-s increments. We then found which lags showed an average regression coefficient that significantly differed from 0 after correction for multiple comparisons across lags (Benjamini and Hochberg, 1995). We first focus on the left hemisphere of the brain (Figure 1A), the hemisphere to which language processing is generally lateralized.

**Figure 1.**
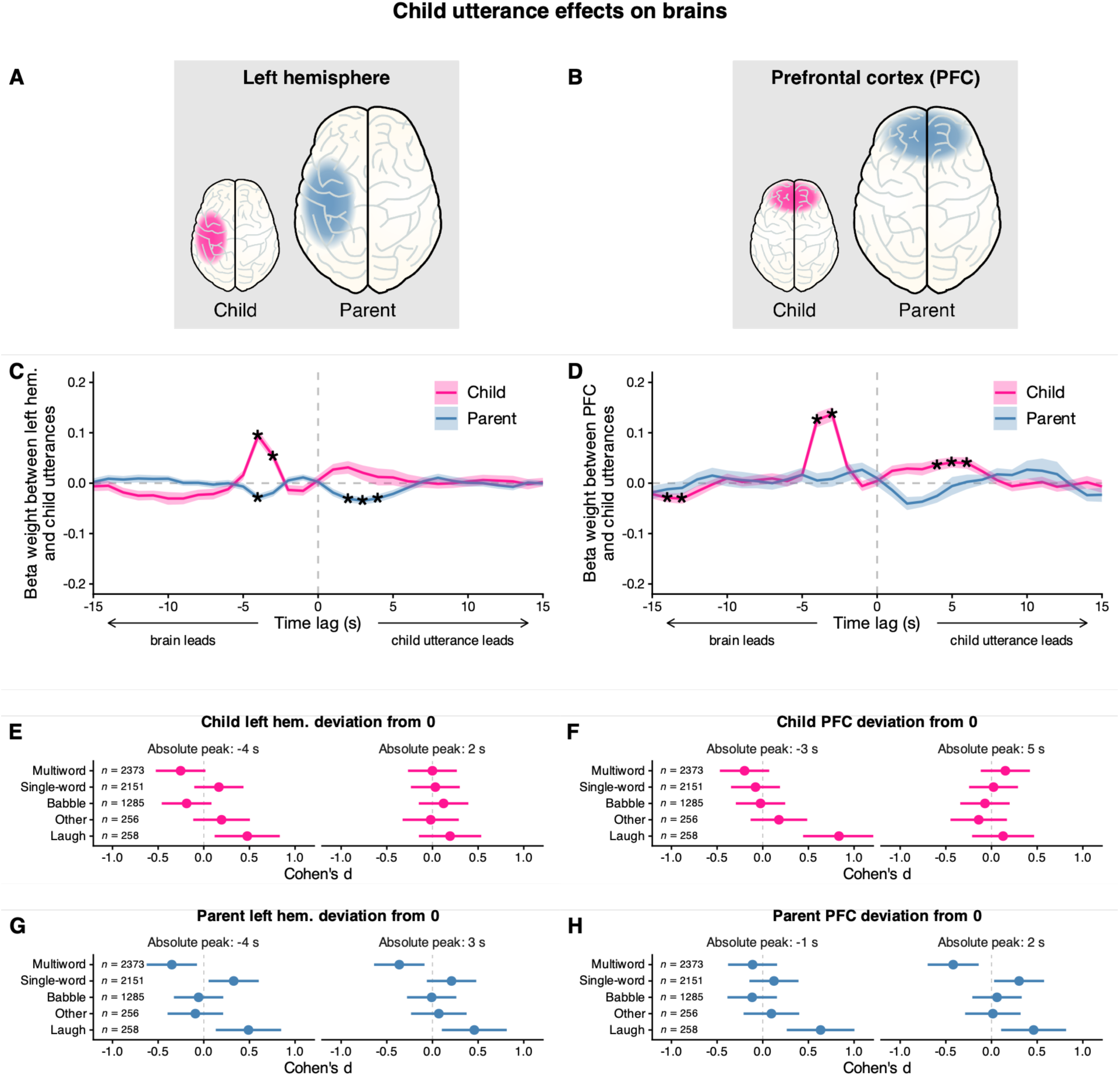
Effect of child utterances on child and parent brains. (**A**,**B**) The regions of interest and brain signals of interest in each column of figures. Results are shown separately for the child in pink and the parent in blue (*n* = 56). Time-lagged relation between child utterances and neural responses in the (**C**) left hemisphere and (**D**) PFC (mean ± 1 SE). Asterisks indicate time lags at which coefficients significantly deviated from 0 after correction for multiple comparisons across lags, * p < .05. Effect size (Cohen’s d ±95% CI) across child utterance types are displayed for moments of peak absolute deviation from 0 for (*left*) brain leading and (*right*) child utterance leading time windows. (**E**) Child left hemisphere effect sizes across types at (*left*) –4 s and (*right*) 2 s. (**F**) Child PFC effect sizes across types at (*left*) –3 s and (*right*) 5 s. (**G**) Parent left hemisphere effect sizes across types at (*left*) –4 s and (*right*) 3 s. (**H**) Parent PFC effect sizes across types at (*left*) –1 s and (*right*) 2 s. These analyses were preregistered.

We found that children’s utterances quickly elicited activity suppression in parents’ left hemisphere activity (Figure 1C). Children’s utterances were reliably followed by their parents’ left hemisphere suppression 2, 3, and 4 s after children produced an utterance. These brain responses were also found in parents’ prefrontal cortex (PFC) and right hemisphere but were most robust in the left hemisphere (Figure 2). Parent left hemisphere suppression was strongest 3 s after children’s utterances (*t*(55) = –4.47, *p* = .0012, Cohen’s *d* = –0.59, 95% CI *d* = [-0.88, – 0.30]) and 42 / 56 parents showed negative brain-behavior coupling at this lag (binomial test, *p* = .0002). This effect held regardless of whether parents verbally responded to their child or not, suggesting that the effect was indeed related to parent listening, not just speaking (Figure 3A). In contrast, random shuffled dyads lagged regressions showed no significant coupling between parents’ left hemisphere and child utterances at any lag (*p*s > .9570). Thus, children’s utterances led to quick left hemisphere suppression in their parents’ brains, which is a known neural signature of speech monitoring (Eliades and Wang, 2008; Harmon et al., 2024; Ozker et al., 2024).

**Figure 2.**
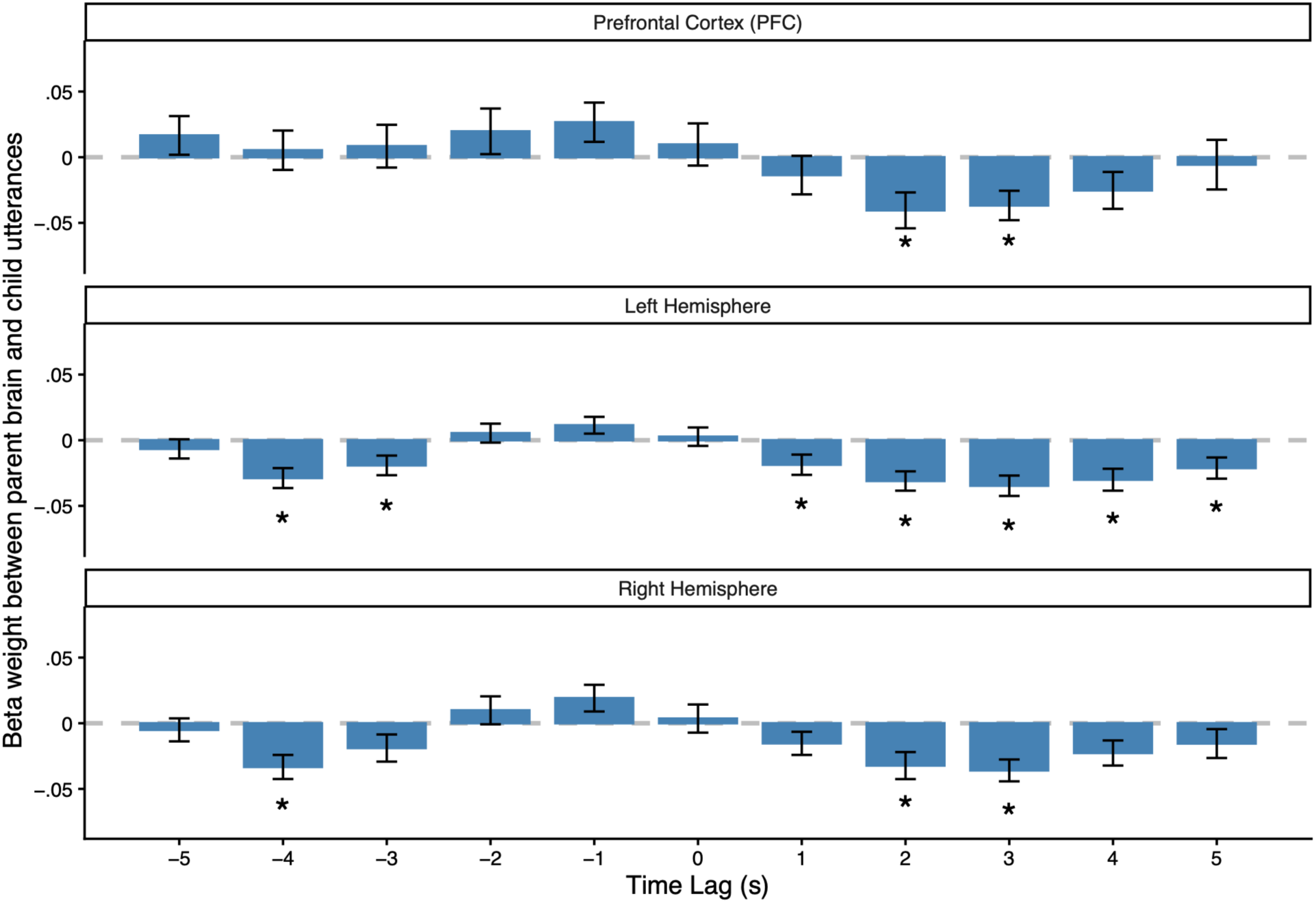
Effect of child utterances on parent prefrontal cortex (PFC), left hemisphere, and right hemisphere. Time-lagged relation between child utterances and neural activity across parent brain regions (mean ± 1 SE, *n* = 56). Children’s utterances elicited a bilaterial listening effect, showing suppression in parents’ left hemisphere, right hemisphere, and to some extent parents’ prefrontal cortex (PFC). We reran the analysis from Figure 1C during the time windows within which our effects were present (–5 to 5 s) across parent brain areas and found evidence that after children vocalized, parents’ PFC, left, and right hemispheres showed suppressed activity 2 and 3 s later. This effect was most consistent in parents’ left hemisphere, showing beta weights significantly below zero 1, 2, 3, 4, and 5 s after children vocalized, correcting for multiple comparisons across lags. Parents’ brains showed significantly different coupling to child utterances after they were produced. Paired-sample comparisons with FDR correction showed significantly different pre-utterance coupling at –1 and –2 s relative to the corresponding post-utterance lags of 1 and 2 s in parents’ PFC, left, and right hemispheres, and a more extended –3 vs. 3 s effect in PFC (*p*s < .0087). This points to a whole-parent-brain activity shift that was promptly organized by children’s early speech production. Asterisks indicate time lags at which coefficients significantly deviated from 0 after correction for multiple comparisons across lags, * p < .05. These analyses were exploratory.

**Figure 3.**
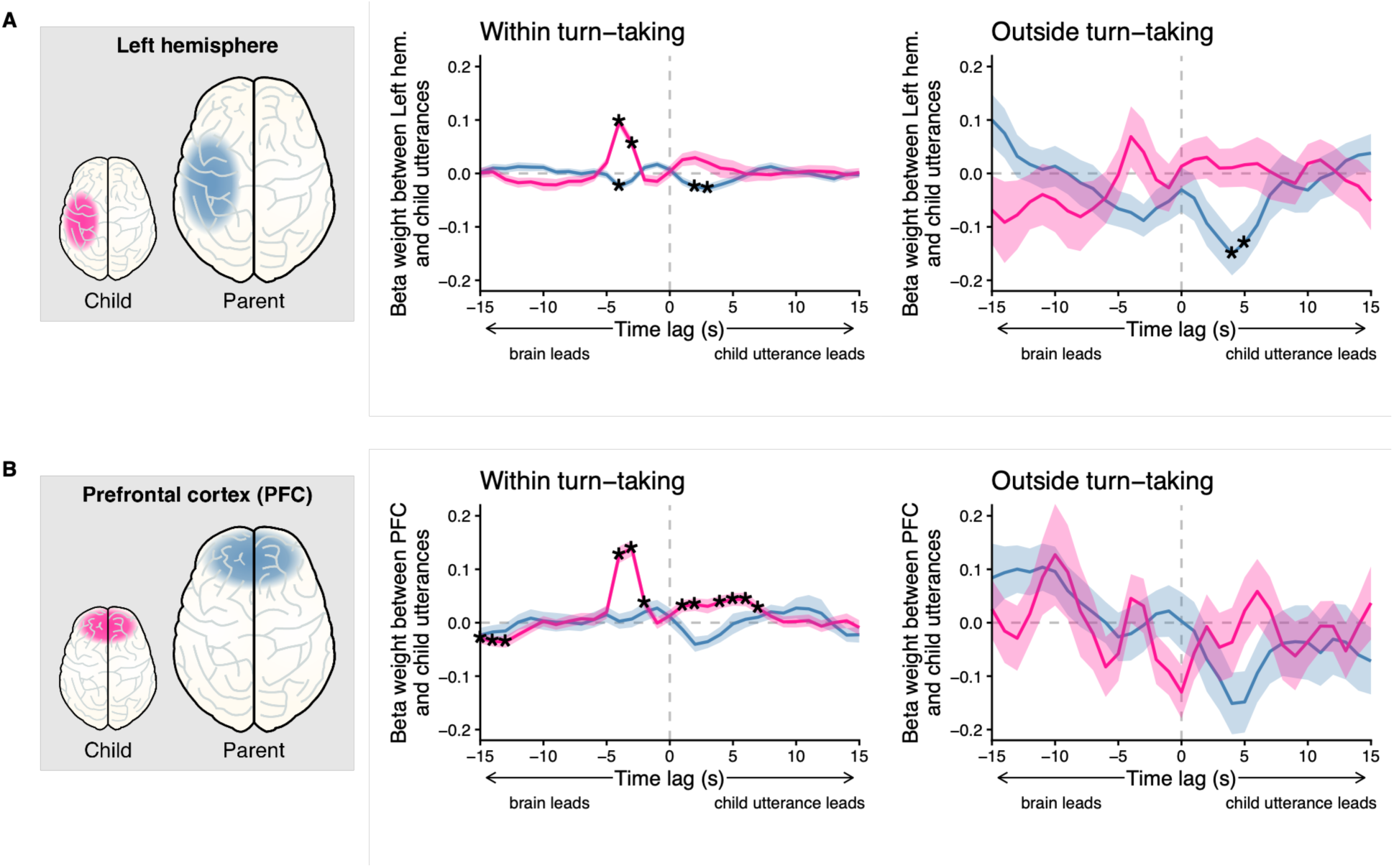
Effect of child utterances on child and parent brain as a function of conversational turn-taking context. (**A**) Lagged brain-behavior regression analyses in the left hemisphere comparing child utterances produced inside of conversational turn-taking with parents vs. outside of turn-taking. Was parent left hemisphere suppression after child vocalizations an effect of parent listening or parent talking? Child vocalizations were very frequently followed by parent verbal responses, with children’s multiword and single word utterances eliciting higher rates of verbal responses (*M* = .82, *SD* = .11) compared to children’s non-speech (i.e., babble, other, and laugh, *M* = .73, *SD* = .14, *p* < .0001). To test listening vs. talking effects, we examined whether parents’ left hemisphere suppression occurred when parents did not verbally respond (i.e., when they produced no conversational turn). While the main temporal dynamics of the brain-child utterance relation happened within turn-taking contexts, parents’ left hemisphere suppression also followed child utterances that were outside of turn-taking. This is consistent with children’s utterances leading to a listening state in parents, not simply a talkative state. (**B**) The same analyses shown in the prefrontal cortex (PFC). Results are shown separately for the child in pink and the parents in blue (mean ± 1 SE, *n* = 56). Asterisks indicate time lags at which coefficients significantly deviated from 0 after correction for multiple comparisons across lags, * p < .05. These analyses were preregistered.

### Child communicative maturity effects in parents’ brains

Did parents’ brains show a sensitivity to the communicative maturity in children’s utterances? We measured whether some child utterance types were more effective than others at suppressing parent left hemisphere activity. When we decomposed the effect size reached by multiple child utterance types during the moment of peak deviation (3 s lag), we found that only children’s multiword utterances showed the predicted suppression effect, with a 95% CI that did not overlap zero (Figure 1G). This pattern was consistent across all three significant post-utterance lags (2, 3, and 4 s), with children’s multiword utterances uniquely and consistently showing negative effect size with 95% CI not overlapping with zero. No other child utterance type showed this pattern. The overall effect after child utterances was negative, consistent with suppression in parent left hemisphere activity (Figure 1C). Importantly, this effect was not related to (a) child age (rho = –.13, *p* = .3100), (b) the total number of child utterances (rho = –.11, *p* = .4160), (c) the total number of child multiword utterances (rho = .02, *p* = .8510), or (d) the duration of child utterances (see *Materials and Methods* for further detail). Together, this suggests that the linguistic complexity of the multiword utterances produced by the child drove the quick suppression of parents’ left hemisphere activity.

To further test the effect of children’s speech maturity on their parents’ brains, we asked 4 independent raters to listen to all of the single– and multiword utterances in our dataset and assess each for how many recognizable English words they heard the child produce. With 4 raters’ recognizability scores (Borjon et al., 2024) we could take an average recognizability per utterance to assess each child’s utterance complexity as a continuous distribution (Figure 4A). We found that parents’ PFC was sensitive to children’s utterance recognizability (Figure 4B). When we partitioned each child’s own utterances into high vs. low recognizability by median split, parents’ PFC showed higher activity 2, 3, 4, and 5 s after lower-recognizability child utterances than higher-recognizability utterances (peak difference at 3 s, *t*(53) = –3.62, *p* = 0.0039, Cohen’s *d* = –.49, 95% CI *d* = [-.76, –.22]). This result is likely to reflect increased effort to understand the less-recognizable content of child speech (Dimitrijevic et al., 2019; Meylan et al., 2023; Vaisberg et al., 2024). Linear models controlling for child age showed that children’s average utterance recognizability was negatively related to their parents’ PFC response to child multiword utterances at 2-5 s lag (*p*s < .0394, Figure 4C). We did not see this sensitivity in parents’ left (*p*s > .9060) or right (*p*s > .8750) hemisphere. Thus, while parents’ brains were generally more sensitive to child utterance complexity, higher-order processing in the PFC may be sensitive to utterances at the upper end of children’s linguistic abilities (Hasson et al., 2008; Peelle, 2018; Sherafati et al., 2022; Wild et al., 2012).

**Figure 4.**
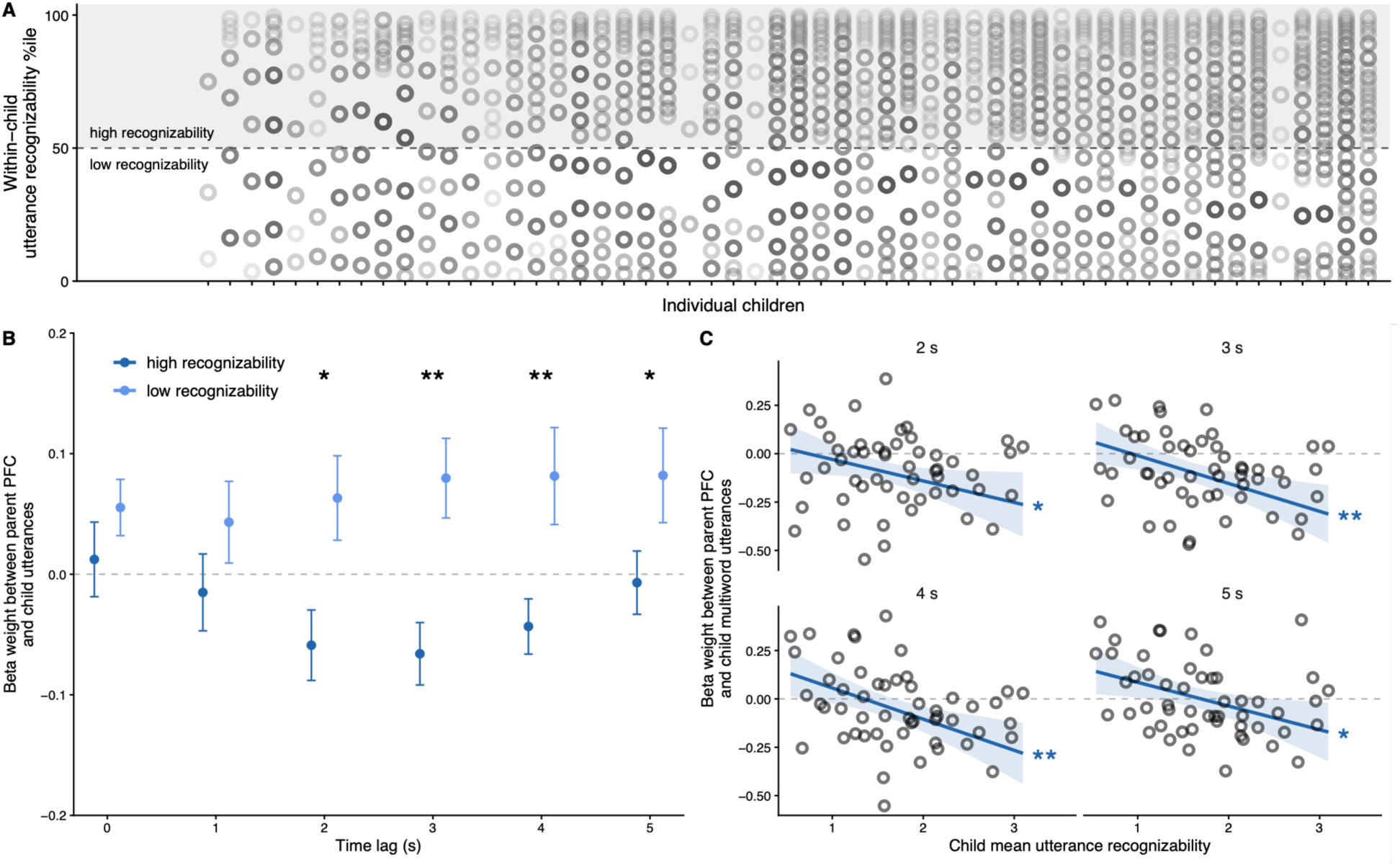
Parent prefrontal cortex (PFC) is sensitive to child utterance recognizability. Children’s highly recognizable utterances elicited suppression in parent PFC relative to low recognizability utterances. This effect is apparent in real-time and at the session level. (**A**) Each child’s utterances were median split by recognizability score into high recognizability and low recognizability. (**B**) High recognizability (dark blue) and low recognizability (light blue) utterances were examined for effects on parents’ brains. Only the parent PFC showed a sensitivity to child utterance recognizability, left and right hemisphere responses were not significant. Parents’ PFC responses 2-5 seconds after low recognizability utterances showed significantly higher brain-behavior coupling than high recognizability utterances (mean ± 1 SE, *n* = 56). Asterisks indicate time lags at which high and low recognizability coefficients significantly differed after correction for multiple comparison. (**C**) The effects of child utterance recognizability were also significant in continuous measures relating children’s overall mean utterance recognizability and parent PFC response to child multiword utterances at 2-5 seconds after their production (shaded bands indicate ± 95% CI, *n* = 56). These analyses were exploratory. * *p* < .05, ** *p* < .01

Parent brain responses may reflect children’s communicative capacities beyond the utterance level, including children’s ability to have coordinated conversation. Consistent with this possibility, children who avoided overlapping speech with their parent elicited a stronger listening effect in parents’ left hemisphere at 2 s lag (*β* = .22, *p* = .0208; see *Materials and Methods*). Thus, parents appear to be sensitive to their child’s communicative maturity, both their speech complexity and conversational skill.

### Children’s brain responses to their own vs. parent speech

We next examined children’s neural sensitivity to the social effects of their *own* speech, which is important for understanding the possibility that parental listening effects are indeed part of a communicative feedback loop with children. We found that children’s own brains (specifically in PFC) were particularly sensitive to moments around their own talk compared to their brain dynamics around parents’ talk. After children vocalized, their PFC showed a reliable increase in activity at 4, 5, and 6 s lags (Figure 1D). Children’s PFC increase peaked 5 s after they spoke (*t*(55) = 3.74, *p* = .0045, Cohen’s *d* = .50, 95% CI *d* = [.215, .783]), with 39/56 children showing positive brain-behavior coupling at this lag (binomial test, *p* = .0045). As shown in Figure 1F, multiword utterances and laughter were numerically highest in driving this PFC increase.

If children are sensitive to the social effects of their vocalizations, we might expect differences in their brain responses after vocalizing inside vs. outside of conversational turn-taking with their parent. To examine whether child utterance effects on their own PFC were sensitive to social context, we analyzed brain-behavior relations inside and outside of conversational turn-taking with parents. Inside of turn-taking bouts, children’s PFC activity significantly increased after vocalizing, showing extended brain-behavior coupling at 1, 2, 4, 5, 6, 7 s after their own utterances (Figure 3B). No timepoints outside of turn-taking were significantly different from zero. Directly comparing children’s PFC response after their own utterances produced inside vs. outside of turn-taking revealed a trending difference at 1 s lag after FDR correction (*p* = .0637), with two-thirds of infants showing this direction of results (binomial test, *p* = .0436). This shows that children’s brains were engaged with the effects of their own utterances in the moments after their own production, particularly inside of conversational turn-taking contexts with their parent.

In contrast to children’s brain responses to their own talk, responses to their parents’ speech were more subtle. Only a single time lag showed significant coupling between children’s PFC and their parents’ speech at –5 s lag (*t*(55) = 3.61, *p* = .0291, Cohen’s *d* = .50, 95% CI *d* = [.21, .79]), Figure 5D). This increase in child PFC mostly anticipated parents’ laughter, and parents’ single-word utterance production 5 s later (Figure 5F).

**Figure 5.**
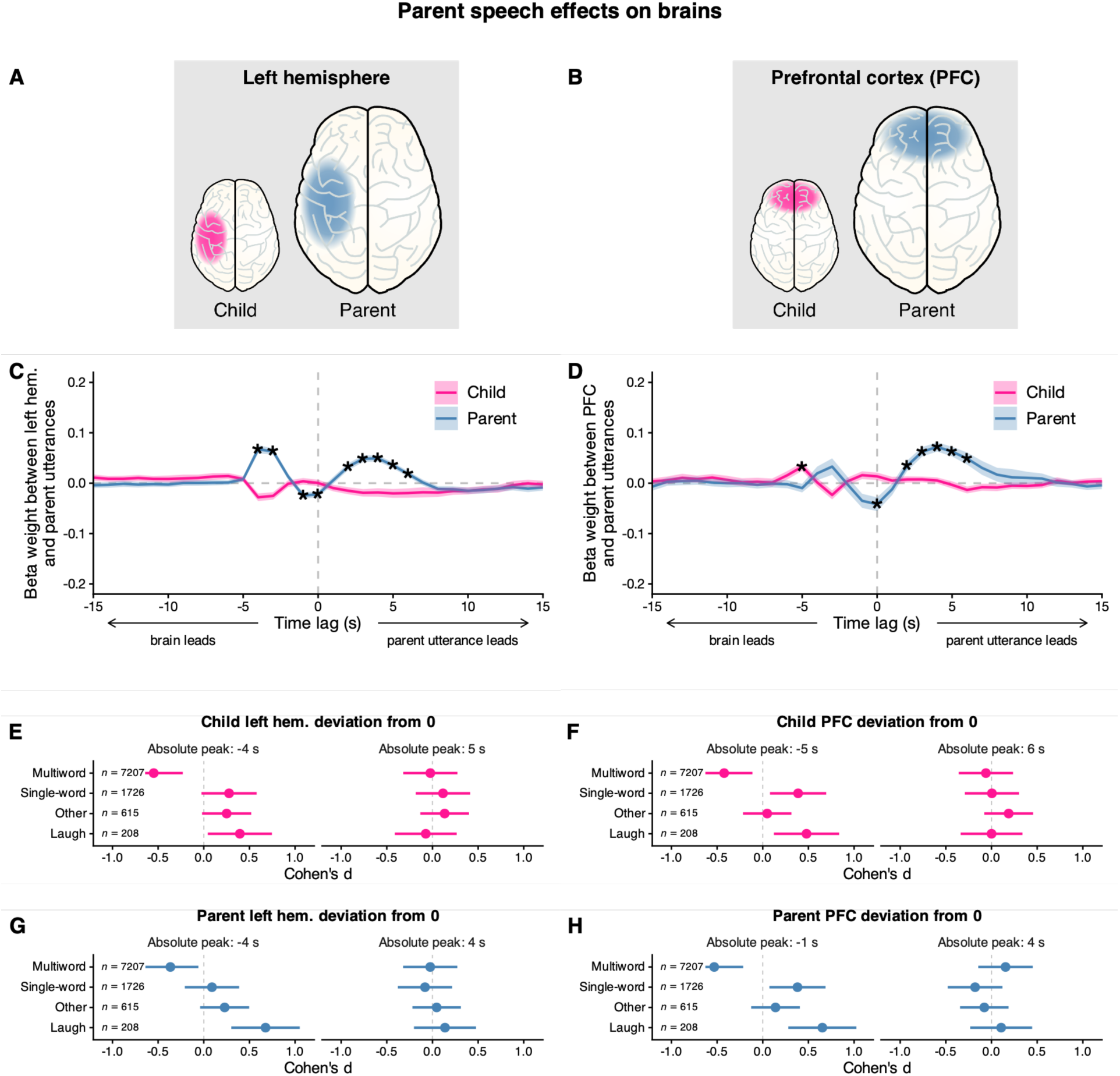
Effects of parent utterances on child and parent brains. (**A**,**B**) The regions of interest and brain signals of interest in each column of figures. Results are shown separately for the child in pink and the parent in blue (*n* = 56). Time-lagged relation between parent utterances and neural responses in the (**C**) left hemisphere and (**D**) PFC (mean ± 1 SE). Asterisks indicate time lags at which coefficients significantly deviated from 0 after correction for multiple comparisons across lags, * p < .05. Effect size (Cohen’s d ±95% CI) across parent utterance types are displayed for moments of peak absolute deviation from 0 for (*left*) brain leading and (*right*) parent utterance leading time windows. (**E**) Child left hemisphere effect sizes across types at (*left*) –4 s and (*right*) 5 s. (**F**) Child PFC effect sizes across types at (*left*) –5 s and (*right*) 6 s. (**G**) Parent left hemisphere effect sizes across types at (*left*) –4 s and (*right*) 4 s. (**H**) Parent PFC effect sizes across types at (*left*) –3 s and (*right*) 4 s. These analyses were preregistered.

To directly compare child brain responses to their own utterances vs. parent utterances, we examined within-subjects comparisons at 0-6 s after production, using FDR correction of *p*-values across lags. We found significantly increased child PFC activity in response to their own utterances at 5 and 6 s lag (*p*s < .0067), with at least 37 out of 56 children showing greater activity after their own utterances at these lags (binomial tests, *p*s < .0222). Additionally, children’s left hemisphere responses to their own utterances were significantly greater compared to parent utterances at 1-3 s lag (*p*s < .0219), with at least 36 out of 56 children showing greater activity after their own utterances at these lags (binomial tests, *p*s < .0440). Thus, parent speech played less of a role in organizing the temporal dynamics of children’s brain activity than children’s own utterance production.

## DISCUSSION

A complete account of experience-dependent development requires a characterization of how developing systems effect change on their immediate environment (Elmlinger et al., 2023; Nikolaus and Fourtassi, 2026; Smith et al., 2018). The field does not currently have a neural account of how children make their parents listen to them. By examining the brain-behavior relations during real-time parent-child interaction, we demonstrate that parents’ brain activity is quickly influenced by their child’s vocalizations. The children in this study were tested during a developmental period marked by regular mixing of immature and mature utterances (McCune and Vihman, 2001). Parents’ neural responses tracked their children’s utterance complexity at both categorical and continuous levels: left hemisphere suppression was driven by children’s multiword utterance types, while parents’ PFC responses were related to children’s speech recognizability. The results additionally show that children’s brains engage with the social consequences of their own vocal productions. Together, these findings capture a neural underpinning of a foundational context of human language learning: communicative exchange between caregivers and children.

Our results are consistent with a fundamental precondition for the idea that feedback from parents drives children to produce increasingly complex vocalizations. Simply getting one’s communication partner to listen may constitute a feedback signal to the learner that guides them toward mature language use (Meylan et al., 2023; Newport et al., 1977; Nikolaus and Fourtassi, 2023). Parents’ brains did not respond uniformly to children’s productions and instead responded selectively to more mature speech, suggesting that children are more likely to receive feedback from parents when they push the boundaries of their communicative skills.

While children obviously do not have perceptual access to their parents’ brain activity, they likely do have access to the behavioral consequences of those brain changes across time and contexts. There may be many instances of communicative success, such as a child saying “uh” and a mother promptly picking them up (Meylan et al., 2023). While our behavioral measures revealed differences in parent response rates to children’s speech vs. non-speech utterances, only our brain measures revealed parents’ sensitivity to the upper end of children’s linguistic capacities. The full suite of features in parental behavior that children could use as feedback is not fully known. Here we use parent brain responses to child utterances as an initial step towards constraining the hypothesis space on this question.

Parents showed robust brain activity suppression following children’s vocalizations during interaction. There was a significant decoupling between child utterances and parents’ left hemisphere activity, indicating that shortly after child talk, parent left hemisphere activity decreased. We additionally found parent left hemisphere suppression inside and outside of conversational turn-taking during the interaction (Figure 3). This auditory suppression after vocalization is a robust cross-species response in animals that communicate acoustically (Eliades and Wang, 2008; Harmon et al., 2024; Ozker et al., 2024), but has not to our knowledge been shown during social interaction. Recent advances suggest that the brain areas (e.g., the left superior temporal sulcus) that show the highest amount of suppression also show the highest amount of activity increases during speech errors, consistent with interpretations of auditory suppression as a speech monitoring signal (Ozker et al., 2024). Our results dovetail with these findings and suggest that parents may exhibit auditory suppression after their children vocalize because they are actively monitoring children’s speech, an interpretation also in line with recent computational models of adults’ child-directed listening (Meylan et al., 2023). If this interpretation is correct, then parents’ suppression should be strongest when children’s productions are most speech-like.

Indeed, parent left hemisphere suppression was primarily driven by children’s multiword utterance production. No other child utterance type elicited parent left hemisphere suppression, and this effect was stable across multiple child utterance durations. Additionally, we further inspected children’s speech utterances for the extent of recognizable words they produced and found that parents’ PFC responded in different ways for highly vs. minimally recognizable child speech. Consistent with prior work on listening effort in the adult brain (Dimitrijevic et al., 2019; Vaisberg et al., 2024), we found that higher-recognizability child utterances elicited decreased parent PFC activity compared to lower-recognizability utterances. While our paradigm was not specifically designed to capture listening effort in parent brains, listening is a likely prerequisite for their brain to differentially respond in-the-moment to differences in children’s speech maturity. These findings hold promise in explaining why children broadly move toward more mature language use, because this is what works in their communication system.

Classic models of communication posit that information transfer is the basic function of communication, and by extension one could assume that the reason children show continued speech development is to increase their capacity for transferring information (Shannon, 1948). However, this premise assumes that children know when they are transferring information to a communication partner. Here we suggest a different hypothesis of why children keep making progress with learning to talk, one with less assumptions of knowledge in the child. Our feedback hypothesis of child-directed listening is that children produce increasingly advanced communication over time because of its potency in communicating successfully. That is, children do not need to know when they are transferring information; instead, they only need to be able to perceive that their communicative attempt somehow influenced their social partner. Children could use the information both from communicative attempts that influence the social partner, and those that fail to do so (Ritwika et al., 2020).

We provide additional support for this hypothesis by showing that children are more engaged in the effects of their own talk, compared to the talk of their parent. Challenges to previous feedback hypotheses focused on rarity of feedback in the linguistic signal of caregiver speech itself (Marcus, 1993). When we broaden the lens to encompass more feedback sources than parents’ child-directed speech alone, it is clear that we need to rethink the idea of feedback being rare. Scaling our findings up to everyday interaction, children are likely to experience feedback almost constantly, based on even subtle multimodal behaviors (or lack of them) in parents. There is evidence that children care about linguistic feedback (Chouinard and Clark, 2003; Elmlinger et al., 2026; Lopez et al., 2020; Newport et al., 1977), and given our findings that their brains are engaged with the effects of their speech during real-time interaction, it seems reasonable that their engagement with feedback on their speech would scale broadly to day-to-day life. We observed clear brain sensitivity in parents to children’s communicative maturity in only 10 minutes of naturalistic interaction. Scaling our findings of child-directed listening to children’s everyday lived experience, the implications are that feedback is ubiquitous for children who produce speech in a socially responsive environment.

While parents’ PFC showed *suppression* after children’s highly recognizable speech, they showed *increased* PFC activity in response to children’s minimally recognizable speech. The former could reflect parents’ inhibition of taking their next conversational turn (Iwaki et al., 2021) as they wait for their child to finish producing speech. The latter is likely to reflect increased effortful cognitive processing (Dimitrijevic et al., 2019; Meylan et al., 2023; Peelle, 2018; Sherafati et al., 2022; Vaisberg et al., 2024; Wild et al., 2012). The timing and selectivity of parent brain response is consistent with previous neuroscience work showing that semantic and discourse content is processed on the order of several seconds, with lower-level information in left hemisphere and higher order in frontal cortex (Lerner et al., 2011).

There are key limits to our study and its implications. First, although the main strength of our design is the prioritization of naturalistic interaction, we therefore do not have the tight control that characterizes distilled lab experimental designs. Future research will need to carry out carefully controlled experiments to further understand parents’ PFC response to children’s high– and low-recognizability utterances. Second, while we have uncovered a neural marker in parents that helps explain children’s progress in learning to talk, there are limits to this claim. For example, when children produce ungrammatical utterances that parents readily and easily understand, children get feedback that their utterance was successful in influencing their communication partner, but they would not receive a cue that improvement to their grammar is called for (Marcus, 1993; Meylan et al., 2023; Nikolaus and Fourtassi, 2023). Future work will need to delineate additional ways that children use their communicative success as a feedback source to guide what to learn next.

## MATERIALS and METHODS

We used dual-brain functional near-infrared spectroscopy (fNIRS) to continuously measure the brains of *N*=56 dyads of toddlers (age ranged from 24 to 43 months, *M* = 31.61) and their parents (Figure 6A) during unstructured play. Each parent-child dyad participated in three four-minute toy play sessions, each with a new set of toys (order counterbalanced). We measured oxygenated hemoglobin concurrently from both parents’ and children’s fNIRS caps with 17 channels on the left hemisphere, 17 on the right hemisphere, 7 channels on the prefrontal cortex (PFC), and time-aligned these signals to child utterances produced in the play session (Figure 6A *Bottom*). For any given frame of neural data (10 Hz), we had information about whether the parent or the child was talking, whether they were in active conversational turn-taking, and what type of utterance they produced. Interactions were approximately 10 minutes in duration (*M* = 9.77, *SD* = 2.03). Across all 56 dyads, we analyzed a total of 6438 child utterances and 9151 parent utterances. On average, children produced 115.0 utterances (*SD* = 37.53), 94% of which were produced in conversational turn-taking bouts with their parent. Parents produced 194.5 utterances on average (*SD* = 48.40), 84% of which were inside of turn-taking bouts with their child. We mainly focused on temporal brain-behavior coupling dynamics because, in order to examine the sensitivity of parents’ brains to the communicative maturity of children’s speech, we need to observe the time course of brain activity before and after children produce vocalizations and early speech. To validate the signal quality of our NIRS recordings, we first examined spatial coupling in true dyads vs random dyads, which replicated brain-to-brain findings from prior studies (Figure 6B; (Piazza et al., 2020; Simony et al., 2016).

**Figure 6.**
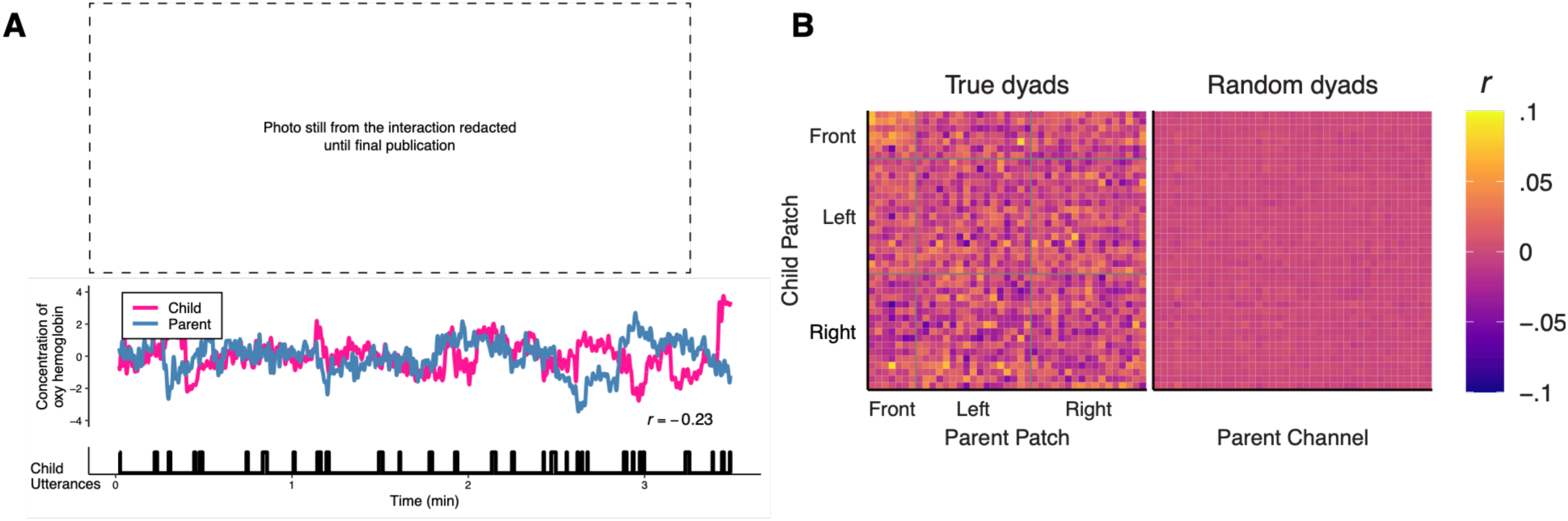
Overview of parent-child dual-brain fNIRS study. (**A**) Example of an interaction between parent and child and the corresponding intersubject correlation (ISC) in one channel pair. The neural timeseries shows concentration of oxyhemoglobin across an excerpt of a 10-minute interaction, separately for the child in pink and the parent in blue. These neural time series unfold alongside real-time utterances produced by the child in black. The ISC between parent and child computed from a homologous left hemisphere channel pair, is shown in the bottom right. (**B**) Spatial extent of parent-child neural coupling. The intersubject correlation matrices show the relation between all parent and child channels, separately for the true dyads (*n* = 56) and randomly-shuffled channel dyads. This illustrates that true parent-child dyads showed spatially coupled brain signals across channels compared to random dyads. To find statistically significant coupling in parent-child channel pairs while accounting for the temporal autocorrelation of the fNIRS signal, we used phase-randomized permutation tests (20,000 surrogate samples) with *p*-values derived from the proportion of randomized correlations that exceeded the observed correlation, with FDR correction for multiple comparison across all channel pairs. Thirty-three channel pairs were significantly coupled under this analysis, most linked parent-child frontal channels (11 frontal-frontal pairs), as well as frontal-lateral and lateral-lateral pairs. This spatial coupling measure (Pearson correlations) shows that we captured significantly coordinated brain-to-brain activity fluctuations during the dyads’ interaction.

### Participants

Fifty-six 24-to 43-month-old children (*M* = 31.60 months, *SD* = 5.56, 34 female), and their parents participated in this study. All children had no history of vision or hearing problems, no known developmental delays, were monolingual (heard above 80% English at home), and were full-term (>37 weeks gestation). In an optional post-session survey, we asked parents demographic questions. Of the parents who completed the survey we observed that children were mostly white (*n* = 36 White, *n* = 5 Asian, and *n* = 9 more than one race) and came from higher-SES households. All mothers had graduated high school, and most had completed college (*n* = 54; 98%) or graduate school (*n* = 38; 69%). Each adult participant was their child’s parent and provided informed consent for their child; no other categories of caregivers or legal guardians were tested in this sample. Photographs from the interaction (Figure S1A) were provided consent for use by families in a follow up email.

Two children were excluded because of technical issues with NIRS data recording. Three children were excluded because they refused to wear the fNIRS cap. Five participants were excluded from behavioral (speech) analysis due to technical issues with the audio/visual recordings or the child vocalized fewer than 5 times. A total of 56 participants provided complete data (behavior and neural) to the present analyses.

### Procedure

We utilized a dual-brain LABNIRS system (Shimadzu Scientific Instruments, Columbia, MD) to simultaneously measure brain activity from both children and their parents over time. We recorded from 41 channels which spanned the cortex of each participant, 7 channels dedicated to their PFC, 17 to the left hemisphere, and 17 to the right hemisphere. The locations of these regions of interest (ROIs) were homologous across child and parent. The center point of the cap was positioned at approximately Cz, thus positioning the caps on the basis of known anatomical landmarks according to the international 10-20 system. Light intensity was recorded in three wavelengths: 780 nm, 805 nm, and 830 nm. Our primary analysis of time-lagged brain-behavior regressions focused on concentrations of oxyhemoglobin (HbO) because prior literature has found greater correlations between fMRI BOLD signals and HbO for naturalistic stimuli (Liu et al., 2017; Piazza et al., 2021). HbO and HbR analyses can be found in our OSF repository.

Parents and their children were welcomed into the lab, asked to sit at a table facing one another, and were equipped with fNIRS caps, first the parent and then the child. Experimenters then explained the task in which parents and children would play with three different sets of toys, which together capture children’s range of experiences with familiar and novel play interactions. One set consisted of familiar toys (*dog*, *spoon*, *banana*), the second consisted of unfamiliar toys (*ferret*, *whisk*, *radish*), and the third consisted of novel toys with novel labels (*gazer*, *cheem*, *tobu*). Each block of play with the individual toy sets lasted 3.5 minutes in duration and after each block ended, an experimenter would introduce the new set of toys. Order of toy set was randomized, except the novel toys were always presented last. Each session was recorded from three different camera angles, a view of the child, a view of the parent and a side view of both interacting. Coders used the side view audiovisual recordings whenever possible (Figure S1A). After participating, families were emailed a $10 Amazon gift card.

### NIRS Preprocessing

All preprocessing steps were done using Homer2 (Huppert et al., 2009). We removed motion artifacts using both moving window and spline interpolation (Lorenzo et al., 2019). We also low-pass-filtered (1 Hz) and high-pass-filtered (0.01 Hz) the signal to remove physiological noise and drift, respectively. Before analysis, the fNIRS data was visually inspected to ensure the quality of data from each channel for each participant (18.9% of channels excluded from the sample). Optical density was then converted to chromophores concentrations using a modified Beer-Lambert Law (Huppert et al., 2009). We downsampled the fNIRS time series to 1Hz. The grand means of neural activity across each ROI for the parent and the child were then calculated separately. Our measurements of brain-behavior coupling were conducted with linear models for each ROI and child utterances. In our control analyses, we randomly reassigned neural time series to the behavioral data of a different dyad. This allowed us to examine the extent of brain-behavior coupling that we might expect by chance alone, following procedures in prior infant-adult fNIRS research (Piazza et al., 2020; Roche et al., 2025).

### Coding of child utterances

Child vocalizations and parents’ speech during the interaction were annotated in full. Parents’ speech was transcribed and children’s vocalizations were annotated for utterance complexity. Utterance boundaries were determined using the ACLEW guidelines (Soderstrom et al., 2021), taking into account prosodic contour, complete clauses, and pause lengths of longer than 1 s in duration. Child utterances were classified as multiword utterances if they contained at least two unique recognizable words in English, a single-word utterance if at least one word was recognizable, a babble if the utterance had at least one clear consonant or vowel produced, a laugh if adult-like laughter was produced, and other if the utterance contained only a vegetative sound (e.g., cough, sneeze or burp). We measured intraclass correlation (ICC) on approximately 15% of the dataset for which child utterances were double-coded. ICC for child utterance category was .92 (multiword utterance = .90, single-word utterance = .63, babble = .91, laugh = .86, other = .00). Parent speech utterance categories were classified in R using the parent speech transcription to count the number of words and detect transcription of laughter.

### Child utterance recognizability

To further assess the linguistic complexity in children’s speech, 4 independent coders counted the number of recognizable English words in all of children’s single– and multiword utterances in the dataset. This allowed us to better understand the neural dynamics surrounding children’s production of linguistic complexity beyond the multiword utterance category. Each child’s speech utterances were rated 4 times for the number of recognizable words it contained. Each utterance was then given a recognizability score corresponding to the average number of recognizable words heard across the four coders (Borjon et al., 2024).

### Child utterance duration control analysis

If the duration of children’s utterances was driving the effect of multiword utterances on parents’ left hemisphere, then equating all utterance types by duration would make the effect of multiword utterances disappear. However, this is not what we observed. By examining only child utterances that were shorter in duration (between –1 *SD* and the mean) and longer in duration (the mean to +1 *SD*), we examined the effect size of multiword utterances relative to other utterance types within a restricted range of utterance durations. In both of these duration ranges, we found that multiword utterances showed the largest negative effect size across utterance types.

### Turn-taking analysis

We categorized speech from parents and children as a function of whether it occurred within or outside of conversational turn-taking using an objective temporal measure. Utterances were considered turns if they occurred within three seconds of the onset of the previous speakers’ utterance (Elmlinger et al., 2025; Nguyen et al., 2022)). This definition allowed for turns to be overlapping speech between the conversation partners, a critical detail to include in developmental turn-taking research (Nguyen et al., 2022).

We quantified overlap as the proportion of each speaker’s utterances that began before the offset of their communication partner’s utterance (Takahashi et al., 2016). In most languages studied, conversation partners avoid overlapping speech with one another (Dingemanse and Liesenfeld, 2022; Stivers et al., 2009), and avoidance of overlap can be measured as the extent to which speakers deviate from overlap that is expected by chance during the interaction (Takahashi et al., 2016). We calculated the expected overlap by chance using a random sampling procedure. We took random samples of timepoints within the play sessions, taking as many samples as there were observed child utterances (*n* = 6438) and calculated the proportion of timepoints that overlapped with parent speech by chance. We repeated this process 1000 times to obtain our estimated average expected overlap by chance value. Our strength of overlap avoidance measure was the difference, per child, between their observed overlap and the overlap expected by chance (Takahashi et al., 2016).

## Acknowledgements

This research was funded by an NIH NICHD F32 fellowship (F32HD116572) to Steven L. Elmlinger, and an NIH NICHD R01 grant (R01HD095912) to Casey Lew-Williams. We thank Aunyae Romeo, Ibrahim Bata and Davis Hobley for their behavioral coding. Thanks to Adele E. Goldberg, Kenneth A. Norman and the members of the Princeton Baby Lab, for comments and discussions on earlier drafts of this manuscript and to all of the participating children and caregivers.

## Notes

### Competing Interest Statement

The authors have declared no competing interest.

